# The serum from critical COVID-19 patients induces proteomic changes in olfactory neuroepithelial cells that resemble post-covid neurological complications

**DOI:** 10.64898/2026.03.09.710460

**Authors:** Lucía Beltrán-Camacho, Santosh D. Bhosale, María Hidalgo-Figueroa, Alejandra Delgado-Sequera, Daniel Sanchez-Morillo, José I. Perez-Revuelta, Cristina Romero López-Alberca, Martin R. Larsen, Rafael Moreno-Luna, Esther Berrocoso, Mª Carmen Duran-Ruiz

**Affiliations:** Biomedicine, Biotechnology and Public Health Department, University of Cadiz, Cádiz, Spain; Biomedical Research and Innovation Institute of Cadiz (INiBICA), Cádiz, Spain; Precision Biomarker Laboratories, Cedars-Sinai Medical Center, Los Angeles, CA, USA; Psychology Department, University of Cádiz, Puerto Real, Spain; Centre for Biomedical Research in Mental Health (CIBERSAM), Institute of Health Carlos III, Madrid, Spain; Automation Engineering, Electronics and Computer Architecture and Networks Department, University of Cadiz, Cádiz, Spain; Department of Mental Health, Jerez de la Frontera University Hospital, Cádiz, Spain; Medicine and Surgery Department, University of Cadiz, Cádiz, Spain; Biochemistry and Molecular Biology Department, University of Southern Denmark, Odense, Denmark; Laboratory of Neuroinflammation, National Paraplegic Hospital, Toledo, Spain; Neuroscience Department, University of Cádiz, Cádiz, Spain

**Keywords:** Olfactory Neuroepithelium Cells, COVID-19, SARS-CoV-2, Neurological Diseases, Psychiatric Disorders, Proteomic

## Abstract

Post-acute sequelae of SARS-CoV-2 infection (PASC), commonly referred to as Long COVID, comprise a constellation of persistent, recurrent, or newly emerging symptoms that may endure for months or years following acute infection. Beyond respiratory impairment, PASC is characterized by a wide spectrum of extrapulmonary manifestations, among which neurological and neuropsychiatric symptoms are highly prevalent. Reported features include olfactory dysfunction with loss of smell and taste, fatigue, neuroinflammation, cognitive and memory impairment, depression, and anxiety, with some symptoms persisting up to one year post-infection.

Despite increasing recognition of these complications, the molecular mechanisms underlying post-COVID neurological sequelae remain poorly defined. In this study, we employed a label-free quantitative (LFQ) proteomics approach to investigate protein alterations in olfactory neuroepithelium–derived stem cells (ONEs), a unique population of neural progenitors located in the olfactory mucosa at the interface between the respiratory system and both the peripheral and central nervous systems. Due to their anatomical exposure and susceptibility to SARS-CoV-2, ONEs represent a highly relevant translational model for exploring virus-associated neurobiological processes.

ONEs derived from healthy donors were incubated with serum from either asymptomatic PCR-positive individuals (AS; n=4) or critically ill hospitalized patients (CR; n=6). Proteomic profiling revealed a distinct differential protein expression pattern in ONEs exposed to CR serum compared with AS serum. Altered pathways were associated with viral infection responses, respiratory and cardiovascular dysfunction, and notably, cerebrovascular and nervous system disorders. These findings highlight the vulnerability of ONEs to systemic factors associated with severe COVID-19 and provide molecular insight into mechanisms potentially contributing to persistent neurological sequelae in PASC.

**GRAPHICAL ABSTRACT:** 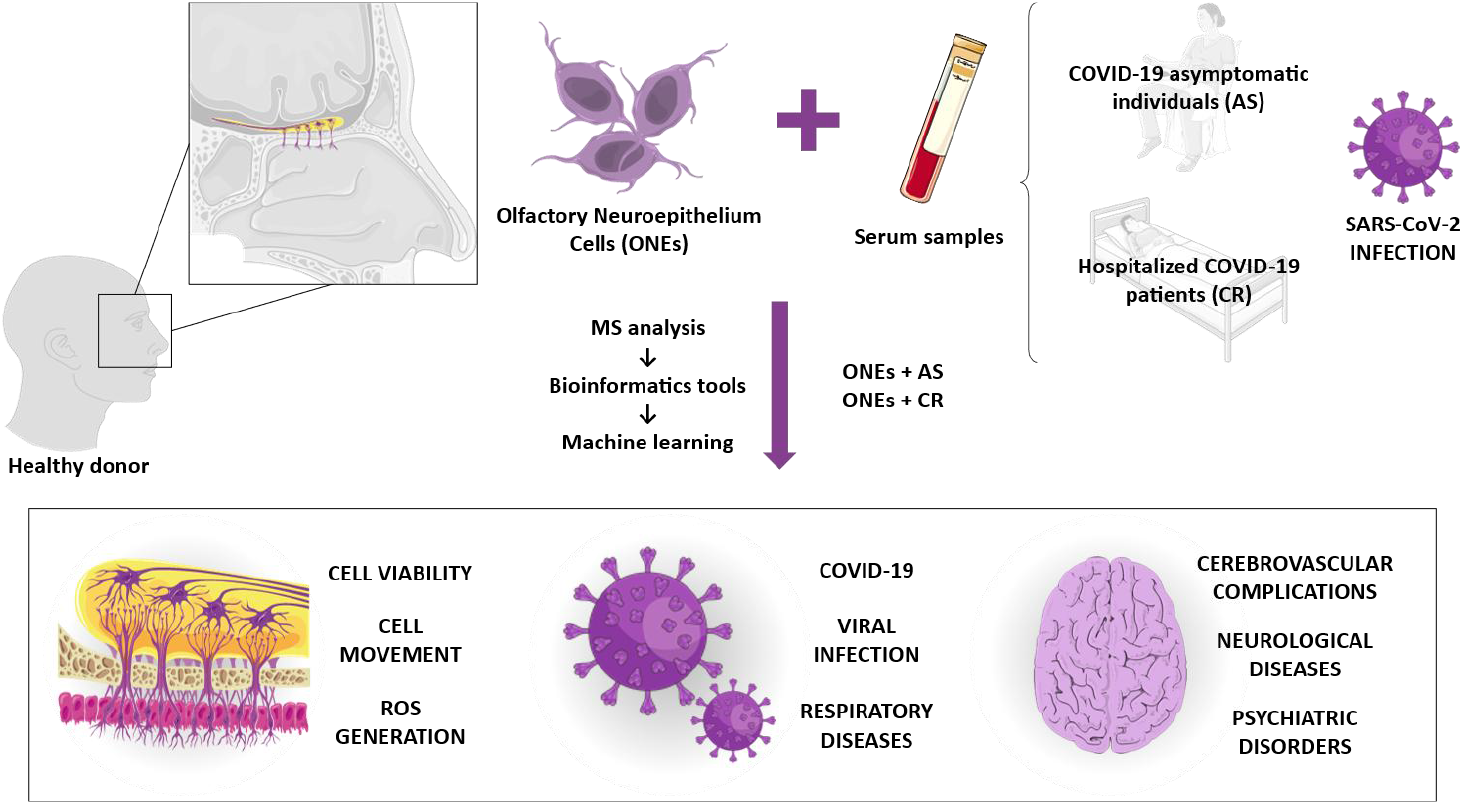

## 1. BACKGROUND

Post-acute COVID-19 sequelae (PASC), commonly referred as long-COVID, encompass a range of physical and mental health issues that persist, emerge, or reappear following an initial SARS-CoV-2 infection. PASC is characterized by a multifaceted array of symptoms that can manifest individually or as a complex combination, posing significant challenges for clinical diagnosis and management [1]. Although SARS-CoV-2 virus primarily targets the epithelial cells of the respiratory tract, causing cough, sore throat, fatigue, fever or upper respiratory complications [2-4], the virus also affects extra-pulmonary organs, including the liver, kidneys, gastrointestinal tract, heart or the brain [5-8]. Post-COVID neurological sequelae involve both peripheral and central nervous system dysfunction. Olfactory impairment, including anosmia and taste disturbances, represents one of the most common features and is likely linked to peripheral nervous system alterations. In addition, patients may experience headaches, neuroinflammation, and cerebrovascular complications. Central nervous system manifestations are also widely reported, including cognitive and memory deficits, language and praxis impairments, reduced psychomotor coordination, and affective disorders such as depression and anxiety [9-13]. Many of these symptoms have been observed even up to 12 months after SARS-CoV-2 infection in hospitalized and intensive care patients, as well as in patients who did not require hospitalization [12]. Notably, more than 30% of COVID-19 severe patients have experienced lasting neurological sequelae or significant changes in mental health [14, 15]. Cognitive disturbances and persistent fatigue represent two of the most burdensome manifestations of long COVID. Notably, these symptoms also serve as key clinical indicators for various neurodegenerative conditions [16]. Unfortunately, some patients have reported neurological disorders which may result in permanent chronic consequences [12]. Indeed, evidence indicates that within 6 months of infection, individuals who contracted COVID-19 face a significantly higher risk of developing Alzheimer’s disease (AD), Parkinson Disease (PD), and multiple sclerosis (MS), compared to those affected by influenza or other respiratory infections [9]. Therefore, COVID-19 might contribute to exacerbate underlying pre-existing conditions or even unmask subclinical neurodegenerative processes [16].

Despite considerable advances in COVID-19 research, the molecular and cellular mechanisms underlying post-COVID-19 nervous system complications remain largely unresolved. It has been postulated that SARS-CoV-2 might access the central nervous system (CNS) via the so-called olfactory pathway [17, 18], potentially spreading from the olfactory epithelium to the olfactory bulb and, in some cases, involving deeper brain structures. Indeed, the virus has been detected in the olfactory mucosa during the acute phase of infection, coinciding with anosmia symptoms [19]. On the other hand, although the olfactory epithelium represents a plausible route for neural invasion, current findings suggest only a limited neuro-invasion through this pathway [1]. Nevertheless, the virus seems to exert a clearly deleterious impact on olfactory cells, leading, among others, to prolonged anosmia [20]. The olfactory mucosa comprises several cell types that have been associated with viral infection, such as support cells and olfactory sensory neurons [19, 21]. In addition, olfactory neuroepithelium cells (ONEs) constitute a significant source of neural progenitors, and a valuable tool for studying the human nervous system [22-24]. Indeed, previous studies have demonstrated that ONEs in vitro resemble molecular and cellular alterations found in patients’ brains [25]. Several studies have suggested that ONEs cells may be an important site of SARS-CoV-2 infection, and subsequent inflammation of these cells may explain the prolonged hyposmia/anosmia [19]. Thus, the use of ONEs constitutes an interesting cellular model for evaluating the long-term effects of SARS-CoV-2 on the nervous system.

To further evaluate the impact of SARS-CoV-2 on ONEs and to better elucidate the neuropathological mechanisms potentially involved, in the current study we applied a proteomic analysis to identify proteins differentially expressed in healthy donor ONEs following ex vivo incubation with the serum from asymptomatic and critical COVID-19 patients. This strategy follows a similar approach to that reported previously [26, 27]. To our knowledge, our findings characterize, for the first time, protein alterations in ONEs that provide significant insights into both the acute nervous system disorders as well as the chronic consequences associated with SARS-CoV-2 infection.

## 2. METHODS

### 2.1. Serum sample acquisition

The study was performed with the serum from asymptomatic individuals (AS, n:4) that tested positive for nasopharyngeal PCR, at the time of blood extraction, recruited at the National Paraplegic Hospital (SESCAM), Toledo, Spain during April-May 2020; and critical COVID-19 donors (CR, n:6) hospitalized at the COVID-19 hospital, Seville, Spain during May 2021. Donor characteristics are shown in Figure 1A-C. Peripheral blood samples were collected using serum separator tubes (SSTTM II advance, BD Vacutainer^®^), centrifuged and stored at −80 ºC, as described [27]. The study was approved by the local Ethics Committee, following Spanish and European Union Regulations and the principles outlined in the Declaration of Helsinki, and donors provided informed consent prior to sample collection. A schematic representation of the work-flow followed in the current study is shown in Figure 1A.

**Figure 1.**
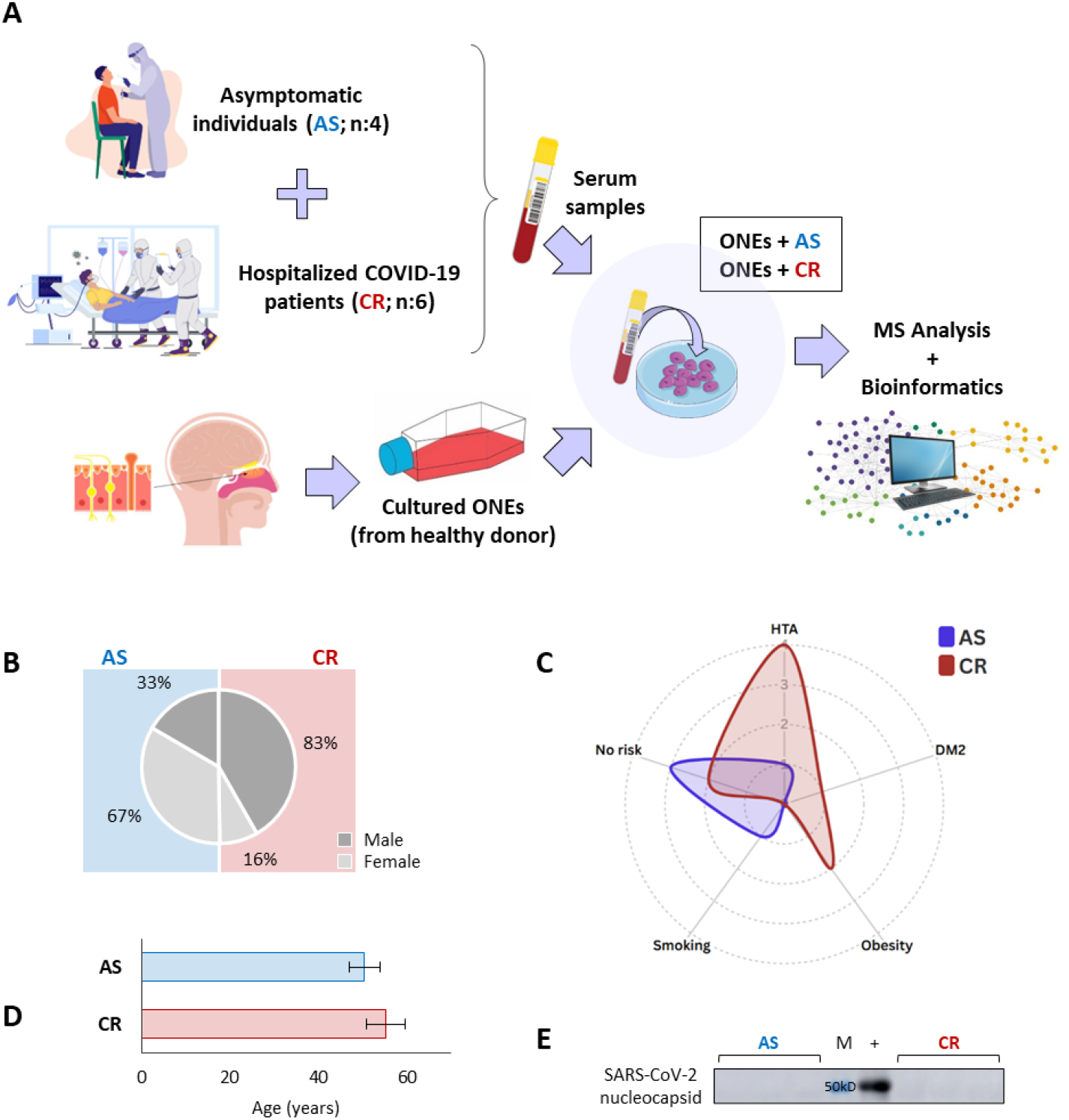
Experimental workflow and study population characteristics. **A)** Schematic representation of the experimental assay: Serum was collected from asymptomatic (AS) and hospitalized critical (CR) COVID-19 patients. Next, ONEs from healthy donors were incubated with the serum samples collected. Proteomic changes were evaluated via a label free quantitative approach, followed by bioinformatics analysis. **B)** Donor gender distribution (percentage). **C)** Prevalence of risk factors across study groups. **D)** Mean age of donors per group. **E)** Western blot analysis for SARS-CoV-2 nucleocapsid protein in serum samples from asymptomatics (AS) and critical (CR) patients. A positive control (+) for SARS-CoV-2 nucleocapside was also included for reference. HTA: arterial hypertension; DM2: diabetes mellitus type 2

### 2.2. Western blot

The presence of SARS-CoV-2 nucleocapsid protein was determined in the serum samples of AS and CR patients by western blot (WB). Proteins were separated by 10% SDS-PAGE, transferred to a membrane and blotted with SARS-CoV-2 nucleocapsid antibody (GeneTex, 135357) diluted at 1:5000. The HRP-conjugated anti-rabbit IgG antibody (GeneTex, 213110-01) was used to detect the primary antibody. A positive control for SARS-CoV-2 nucleocapsid was included to corroborate the presence or absence of this protein in our samples.

### 2.3. ONEs isolation and culture

ONE cells were obtained from a healthy subject, without any diagnostic criteria according to the DSM-5 manual for any severe mental disorder and with no SARS-CoV-2 infection, tested by both nasopharyngeal PCR and blood antibodies at the time of sample collection. Cell culture was performed as previously described [25, 28]. Briefly, cells were exfoliated from the nasal cavity using specific brushes to obtain samples from the lower and middle turbinate through a circular movement. Cells were grown at 37 °C with 5% CO_2_ in Dulbecco’s Modified Eagle Medium/Ham F-12 (DMEM/F12) (GibcoBRL) supplemented with 10% fetal bovine serum (FBS) (GibcoBRL), 2% GlutaMAX 100X (GibcoBRL) and 0.2% primocin (Invitrogen). When 80% confluency was reached, cells were detached with 0.25% trypsin-EDTA (GibcoBRL), replated in 75 cm^2^ flasks and cultured in the same supplemented DMEM/F12 (GibcoBRL). Ethical considerations for the neuroepithelial samples followed the same procedure described in section 2.1.

### 2.4. ONEs incubation *ex vivo* with patients’ serum

ONEs (20*10^5^ cells), in passage 6, were seeded in 12 well plates with DMEM medium plus 10% FBS and 1% antibiotic-antimycotic and incubated 24 h (37 ºC, 5% CO_2_) to settle down. Next, the medium was discarded, cells were washed with PBS 1X to eliminate any remaining traces of FBS from the initial conditioned media, and then incubated for another 24 h with DMEM medium containing 10% serum of the asymptomatic (AS) or critical (CR) groups, as described previously [27, 29]. Cells were collected using 0.25% Trypsin-EDTA (GibcoBRL), centrifuged and washed once with PBS 1X.

### 2.5. Proteomic analysis

The proteome changes of ONEs in response to the incubation with the serum samples from asymptomatic individuals (ONEs+AS, n:4) and critical patients (ONEs+CR, n:6) were analyzed using tandem mass spectrometry (LC-MS/MS) and label free quantitative (LFQ) analysis. Briefly, cell pellets were resuspended in lysis buffer (8M urea plus protease inhibitors; 04693132001, Roche) for protein extraction, and protein quantification was performed with the Qubit Fluorometric system (ThermoFisher Scientific). Next, 50 µg of proteins in 8M urea per sample were reduced (10 mM Dithiothreitol), alkylated (50 mM Iodoacetamide), and diluted four times with 50 mM ammonium bicarbonate for further digestion with Trypsin/LysC (V5073; Promega) (enzyme/substrate ratio 1:50), at 37 °C, overnight. Finally, the digestion was quenched with 1% formic acid (FA) before peptide purification with C18 micro-columns, as described [30], and eluates were dried with a speed-vac system. The dried peptides were reconstituted in 0.1% FA and a NanoDrop (DeNovix, DS-11 Spectrophotometer) was used to estimate the peptide concentration. An amount equivalent to 200 ng of peptides were injected on Aurora Series UHPLC emitter column (250 mm X 75 µm id, 1.6 µm C18) from IonOpticks connected to Easy nLC (Thermo Scientific) interfaced with a timsTOF Pro mass spectrometer (Bruker Daltonics) using the same parameters described in [26]. The instrument was operated in diaPASEF mode via Captive nano-electrospray source (Bruker Daltonics) at 1400 V using an accumulation and ramp time of 100 ms. A 25 m/z precursor isolation width was used to cover 400 to 1200 m/z, covering an ion mobility range /1/K_0_) from 0.60 to 1.60 V.s/cm^2^ [31].

### 2.6. Data processing and statistics

diaPASEF files were analyzed using Spectronaut (v 15.2.210819.50606, Biognosys AG) in directDIA™ mode with default settings. The Spectronaut output containing normalized protein intensities values were then exported into a tabular format for further analysis in the Perseus software [32]. Briefly, protein intensity values were log_2_ transformed and samples were categorically annotated to define the conditions. A t-test differential expression analysis was used with a permutation-based FDR calculation. Proteins were considered as differentially expressed between the groups when p-values < 0.05 and log_2_ foldchange (FC) > 1 (up-regulated) or < −1 (down-regulated). These changes were confirmed afterwards with GraphPad Prism 9 software. Data were presented with box and plots graphs representing median, min and max value and showing all points. The differentially expressed proteins (DEPs) identified between both groups, based on p-values and FC criteria, served as input for STRING and Ingenuity Pathway Analysis (IPA) analysis. STRING database was used to create the protein-protein interaction network, and functional annotation of proteins was performed with IPA. In addition, receiver operating characteristic (ROC) curves were generated for DEPs by plotting sensitivity (%) against 100%-specificity (%), indicating the area under the curve (AUC) and 95% confidence intervals.

## 3. RESULTS

### 3.1. Study population

Two main groups were evaluated in this study: A first group of PCR+, asymptomatic individuals (AS, n:4), and a critical group of patients hospitalized due to serious conditions (CR, n:6). All participants were positive for SARS-CoV-2 nasopharyngeal PCR at the time of blood extraction. Given the early pandemic timeframe, none of the asymptomatic subjects were vaccinated, and only one patient in the critical group had received an initial vaccine dose. (April-May 2020). The mean age of the AS group was 50.38 years, predominantly female (63%), while critical patients were slightly older (55.33 years) and mostly men (83%). The leading risk factor identified among the critical patients recruited was hypertension (HTA) (Figure 1B-D). Finally, the results from WB analysis of the SARS-CoV-2 nucleocapsid protein did not show any trace of this protein in the serum samples of either AS or CR patients (Figure 1E).

### 3.2. The serum of critical COVID-19 patients promotes a differential protein expression profile in ONEs

A total of 4302 proteins were identified in ONEs following exposure to the serum from either critical (ONEs+CR) or asymptomatic (ONEs+AS) COVID-19 patients. The principal component analysis (PCA) revealed a clear segregation between the ONEs+AS and ONEs+CR groups, a finding further supported by hierarchical cluster (Figure 2A-B). Besides, LFQ analysis reported a total of 1649 differentially expressed proteins (DEPs, p-value < 0.05) between both groups. Among these, 86 proteins were significantly up-regulated (fold-change>1) and 113 were down-regulated (fold-change < 1) in ONEs+CR cells compared to ONEs+AS (Figure 2C). The potential interactions found among these DEPs are illustrated in the protein-protein interaction network, Figure 2D. Detailed protein identification and quantitation data are available in Supplementary Table S1.

**Figure 2.**
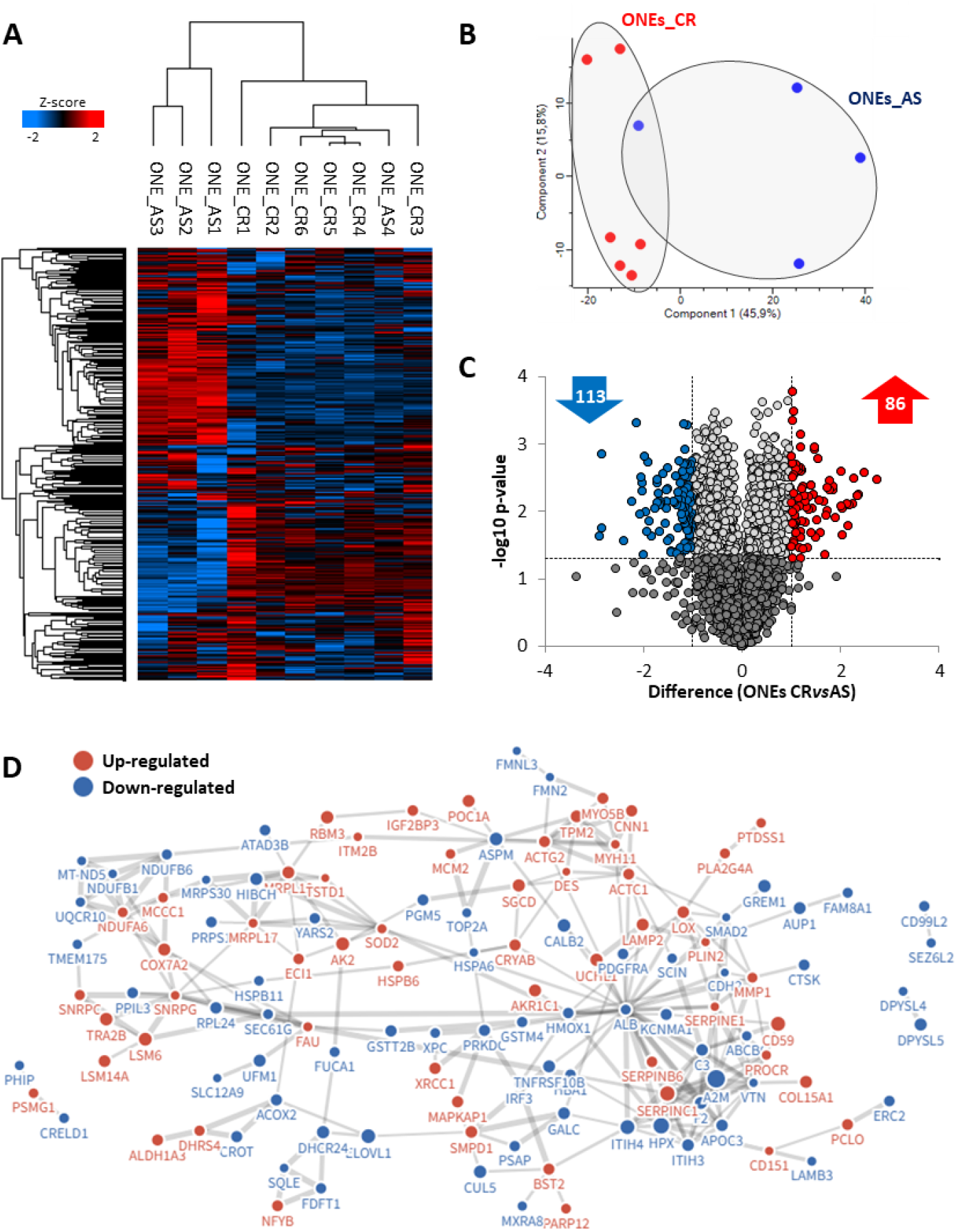
Proteomic profile of ONEs in response to critical (ONEs+CR) and asymptomatic (ONEs+AS) COVID-19 patients’ serum. **A)** Unsupervised hierarchical clustering of the differentially expressed proteins (DEPs). **B)** Principal component analysis (PCA) showing the separation between ONEs+CR and ONEs+AS groups based on their proteomic profiles. **C)** The volcano plot illustrates the distribution of the DEPs, with the number of proteins up-(red) and down-(blue) proteins indicated as well (-Log_10_ p-value >1.5, difference ±1). **D)** Protein-protein interactions network representing potential functional associations among the proteins altered in ONEs+CR *vs* ONEs*+*AS.

### 3.3. The serum of critical COVID-19 patients induces protein changes in ONE cells related to viral infection and respiratory diseases

Functional enrichment analysis of the DEPs identified between ONE+CR and ONEs+AS cells revealed a profound impact of the serum factors from CR patients on ONEs functionality. These alterations primarily affected biological processes related to cell motility and survival, among others (Table 1). Furthermore, canonical pathway analysis highlighted a dysregulation of proteins associated with acute phase response signaling (A2M, ALB, C3, F2, HMOX1, HPX, ITIH3, ITIH4, SERPINE1, SOD2), mitochondrial dysfunction (ATP5MF, COX7A2, MT-ND5, NDUFA6, NDUFB1, NDUFB6, SOD2, UQCR10), coagulation (A2M, F2, SERPINC1, SERPINE1) or complement system (C3, CD59) processes. Moreover, specific pathways related to coronavirus pathogenesis were also reported (Figure 3A). Indeed, regarding the clinical manifestations of severe COVID-19, many altered proteins were closely associated with respiratory pathologies, including pneumonia (ALB, C3, F2, KCNMA1, PDGFRA, SERPINC1, TOP2A) and acute respiratory distress syndrome (ALB, ATP2C1, C3, F2, SERPINC1, SERPINE1), among others (Figure 3B). Furthermore, many of the DEPs identified have been previously linked to viral infection (C3, CRYAB, DPYSL5, F2, IRF3, MT1X, MXRA8, PDGFRA, PTDSS1, RNH1, SNRPC, TMEM63A) or specifically to COVID-19 (ALB, BST2, C3, FAU, F2, ITIH3, ITIH4, LAMP2, MECP2, MMP1, PCLO, PDGFRA, PLA2G4A, PTK7, SERPINC1, SERPINE1, SEZ6L2, SRR, VTN) (Figure 3B-C).

**Table 1.** Functional classification of DEPs in ONEs after CR serum *vs* AS serum incubation. Protein classification based on biomedical literature and integrated databases, performed with the IPA software. The table shows the most probable functions in which the proteins of interest are involved including functions, p-value, activation z-score, molecules (protein names) and number of proteins.

| Functions | -log10<br>(p-value) | Activation<br>z-score | Molecules | Proteins |
| --- | --- | --- | --- | --- |
| Angiogenesis | 3.519 | 0.046 | C3, CD151, CDH2, CRYAB, CUL5, F2, HMOX1, HSPB6, KCNMA1, LOX, MECP2, NAXE, NCKIPSD, PROCR, RNH1, SERPINC1, SERPINE1, VTN | 18 |
| Vasculogenesis | 2.375 | -0.901 | C3, CD151, CDH2, CRYAB, CUL5, F2, HMOX1, KCNMA1, NCKIPSD, PROCR, SERPINC1, SERPINE1, VTN | 13 |
| Apoptosis | 5.777 | -0.517 | ALB, C3, CD59, CDH2, CRIP1, CRYAB, CUL5, DAP, DHCR24, F2, FMN2, GREM1, H1-2, HIGD2A, HMOX1, IRF3, KCNMA1, LAMP2, LCMT1, MCM2, MRPS30, PDGFRA, PEA15, PLIN2, PRKDC, PROCR, PRPS1, RBM3, SCIN, SEC61G, SERPINB9, SERPINC1, SERPINE1, SMAD2, SMPD1, SOD2, TNFRSF10B, TOP2A, UCHL1, VAV3, VTN, XPC, XRCC1 | 43 |
| Cell survival | 3.539 | -1.231 | ABCB6, ALB, C3, CALB2, CD151, CDH2, DHCR24, F2, HMOX1, HSPA6, LAMP2, MCM2, PDGFRA, PLCD3, PLIN2, PRKDC, RASSF8, SERPINE1, SLC16A1, SQLE, TNFRSF10B, TOP2A, UFM1, XPC, XRCC1 | 25 |
| Cell viability | 3.495 | -0.978 | ABCB6, ALB, C3, CALB2, CD151, CDH2, DHCR24, F2, HMOX1, HSPA6, LAMP2, MCM2, PDGFRA, PLCD3, PLIN2, PRKDC, RASSF8, SERPINE1, SLC16A1, SQLE, TNFRSF10B, TOP2A, UFM1, XRCC1 | 24 |
| Necrosis | 3.708 | -1.407 | ALB, APOC3, ATP2C1, C3, CD59, CRYAB, DHCR24, F2, FAU, FDFIT1, HMOX1, IRF3, ITM2B, LAMP2, MCM2, MT1X, PDGFRA, PEA15, PLA2G4A, PRKDC, PROCR, PRPS1, RBM3, SCIN, SEC61G, SERPINC1, SERPINE1, SLC16A1, SMAD2, SMPD1, SOD2, SQLE, TNFRSF10B, TOP2A, UCHL1, VAV3, VTN, XPC, XRCC1 | 39 |
| Autophagy | 2.063 | 1.005 | BST2, DAP, LAMP2, PRKDC, PSAP, SMAD2, SMPD1, SOD2, TNFRSF10B, UCHL1 | 10 |
| Cell spreading | 2.693 | -1.067 | ALB, C3, CRYAB, SERPINE1, SMPD1, VTN | 6 |
| Activation of cells | 2.254 | -1.810 | ALB, C3, CRYAB, F2, MT-ND5, PDGFRA, PROCR, RASSF8, TNFRSF10B, TOP2A, VAV3, VTN | 12 |
| Endocytosis | 2.101 | 0.409 | APOC3, C3, CD151, ELOVL1, HMOX1, LCMT1, SERPINE1, SHOC2, SMPD1, VTN | 10 |
| Cell movement | 3.285 | -0.660 | A2M, ACTC1, ALB, ALDH1A3, BPNT2, BST2, C3, CD151, CDH2, CRYAB, F2, FMN2, FMNL3, HMOX1, IGF2BP3, IRF3, KCNMA1, LOX, MECP2, MMP1, PDGFRA, PHIP, PLPP3, PROCR, PTK7, SERPINC1, SERPINE1, SHOC2, SMAD2, SMPD1, SOD2, TPM2, VAV3, VTN | 34 |
| Invasion of cells | 2.987 | -0.682 | BPNT2, CD151, CDH2, CTSK, DPYSL4, F2, FMN2, FMNL3, HMOX1, IGF2BP3, KCNMA1, LAMB3, LOX, MBD2, MMP1, PDGFRA, PEA15, PHIP, PTK7, SERPINE1, SMAD2, SMPD1, SOD2, VTN | 24 |
| Migration of cells | 3.772 | -0.847 | A2M, ALB, ALDH1A3, BPNT2, BST2, C3, CD151, CDH2, CRYAB, F2, FMN2, FMNL3, HMOX1, IGF2BP3, IRF3, KCNMA1, LOX, MECP2, MMP1, PDGFRA, PHIP, PLPP3, PROCR, PTK7, SERPINC1, SERPINE1, SHOC2, SMAD2, SMPD1, SOD2, TPM2, VAV3, VTN | 33 |
| DNA damage | 3.096 | -1.664 | ACTC1, PRKDC, RNH1, SOD2, TOP2A, XRCC1 | 6 |
| Generation of reactive oxygen species | 2.636 | -1.408 | ALB, CRYAB, F2, ITM2B, SOD2 | 5 |
| Hemostasis | 3.202 | -1.411 | C3, CD59, F2, PDGFRA, PROCR, SERPINE1, VAV3 | 7 |
| Synthesis of fatty acid | 2.402 | -0.248 | ALB, APOC3, ELOVL1, F2, GSTM4, PLA2G4A, VTN | 7 |
| Synthesis of lipid | 4.269 | -0.468 | ALB, ALDH1A3, APOC3, C3, DHCR24, ELOVL1, F2, FDFIT1, FMO3, GSTM4, PEDS1, PLA2G4A, PLPP3, PTDSS1, SMPD1, SQLE, TRA2B, VTN | 18 |

**Figure 3.**
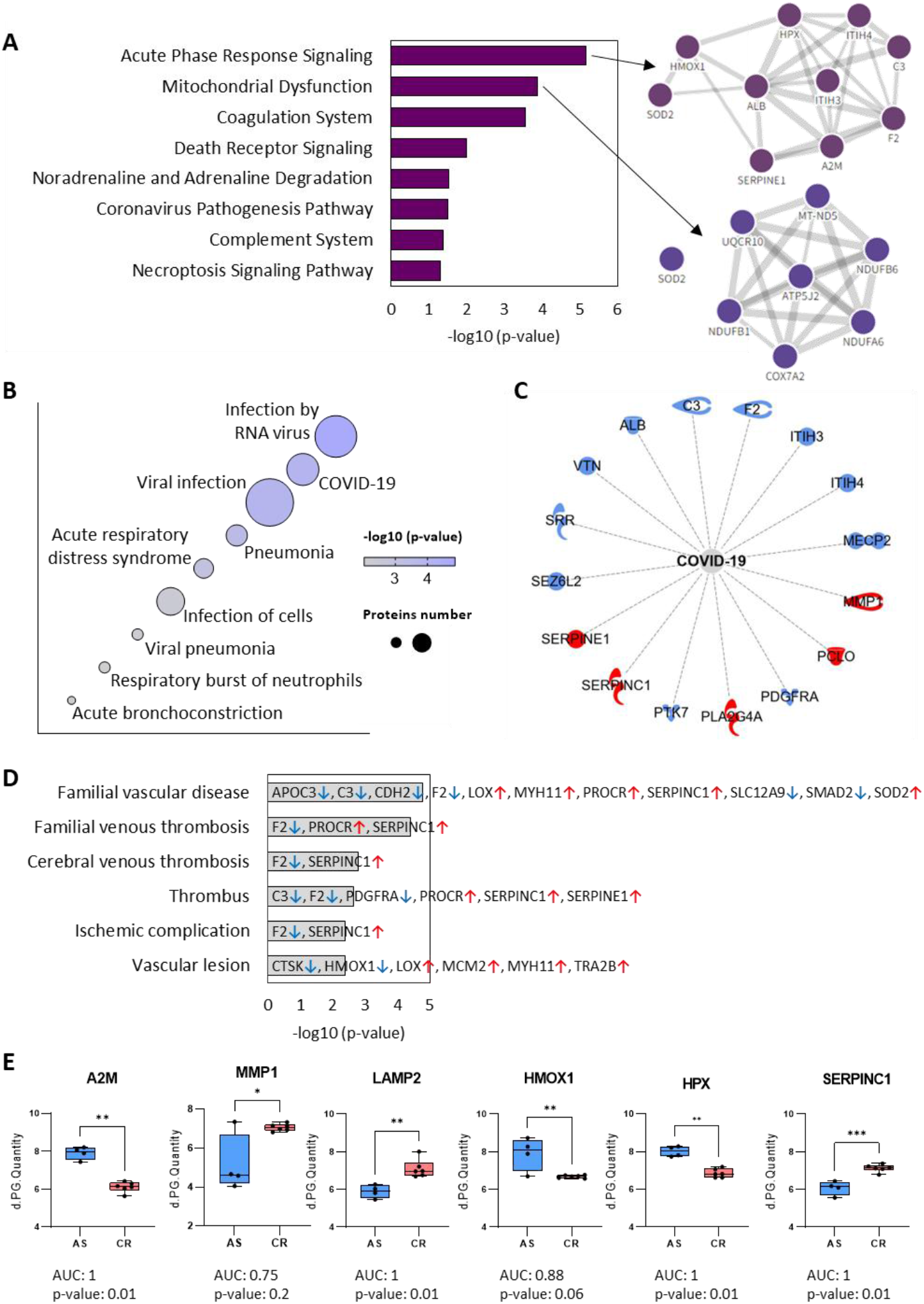
Functional enrichment analysis of differentially expressed proteins (DEPs) in ONEs+ CR *vs* ONE+AS. **A)** Canonical pathways significantly associated with the deregulated proteins. **B)** Functional classification of DEPs linkedto infectious, inflammatory and respiratory diseases. **C)** IPA functional network illustrating the interactions between proteins correlated with COVID-19. **D**) Functional classification of DEPs related associated with cerebrovascular complications. **E)** Graphical representation of LFQ intensities for some of the DEPs identified. Data are presented as box-plots, with the median, minimum and maximum values represented, including single data points as well. Differences were considered significant when *p-value < 0.05. **p-value < 0.01. Data from receiver operating characteristic (ROC) analysis with area under curve (AUC) is also indicated.

Beyond respiratory impairment, and in agreement with previous studies [26], the incubation *ex-vivo* of healthy ONEs with the serum factors from CR patients triggered changes in proteins linked to cardiovascular complications associated with SARS-CoV-2 infection (Figure 3D). For instance, proteins like antithrombin-III (SERPINC1) or prothrombin (F2), were found up- and down-regulated respectively in ONE+CR *vs* ONE+AS cells. Together with other dysregulated proteins, these changes are linked to thrombus formation (C3, F2, PDGFRA, PROCR, SERPINC1, SERPINE1), vascular diseases (APOC3, C3, CDH2, F2, LOX, MYH11, PROCR, SERPINC1, SLC12A9, SMAD2, SOD2), or ischemic complications (F2, SERPINC1). A representative selection of the protein changes identified are shown in figure 3E. Collectively, these findings demonstrate that the serum of critical COVID-19 patients induces a comprehensive proteomic reprogramming in ONEs that mirrors the systemic respiratory and cardiovascular damage observed in these patients.

### 3.5. The serum of critical patients induces proteins changes associated to post-COVID neurological complications and psychiatric diseases

Functional enrichment analysis revealed a significant representation of proteins linked to nervous system diseases in ONE+CR cells (Table 2, Figure 4A-B), including dementia, familial encephalopathy, neurological disorder or familial psychiatric disease. Among these, several proteins up-regulated such as SOD2 and MOCOS have been linked to redox imbalance and neurodegeneration [33-35]. Furthermore, an additional targeted literature search identified further dysregulated proteins like PCLO or BPNT2 (up) and SRR (down-regulated), associated with autism, depression, bipolar disorder or schizophrenia [36-39]. Finally, a significant down regulation of proteins related to neurogenesis, synaptic maintenance or structural integrity (SMAD2, Sez612, NTM, MECP2) [40, 41], was also reported in ONE+CR cells. Altogether, these proteomic alterations seen in ONEs exposed to the serum factors from CR patients are representative of changes in neural plasticity and cognitive health (Figure 4D).

**Table 2.** Nervous system diseases related with DEPs in ONEs after CR serum vs AS serum incubation. Protein classification based on biomedical literature and integrated databases by the IPA software. The table shows the most probable neurological and psychiatric disorders in which the proteins of interest are involved including diseases annotation, −log10 (p-value), protein names and number of proteins.

| Diseases annotation | -log10 (p-value) | Molecules | Proteins |
| --- | --- | --- | --- |
| **Familial neurological disorder** | 6.228 | A2M, ALB, ALDH1A3, ASPM, C3, CD59, CDH2, COL15A1, CRYAB, DHCR24, ELOVL1, EPB41L1, FAM136A, FDFT1, FMN2, FUCA1, GALC, GSTM4, H1-4, HIBCH, HMOX1, IRF3, ITIH4, ITM2B, KCNMA1, LAMB3, MCM2, MECP2, MED17, MMP1, MT-ND5, MT1X, MYH11, NAXE, NDUFB1, NDUFB6, PCLO, PHIP, PRKDC, PRPS1, PSAP, SERPINB6, SERPINC1, SERPINE1, SHOC2, SLC16A1, SMPD1, SOD2, TMEM63A, UCHL1, UFM1, UQCR10, XRCC1, YARS2, YIPF5 | 55 |
| **Progressive neurological disorder** | 5.857 | A2M, ACTC1, ALB, APOC3, BST2, C3, CNN1, CRYAB, DHCR24, ELOVL1, F2, FDFT1, FMN2, FUCA1, GALC, HDAC7, HMOX1, HPX, HSPA6, KCNMA1, MED17, MMP1, MT-ND5, MYH11, NAXE, NOM1, PDGFRA, PIP4P2, PRKDC, PSAP, PTDSS1, SERPINC1, SERPINE1, SMPD1, SOD2, TOP2A, UCHL1 | 37 |
| **Dementia** | 5.323 | A2M, ALB, APOC3, C3, CNN1, DHCR24, F2, FDFT1, FMN2, HMOX1, HPX, HSPA6, ITM2B, MMP1, MT-ND5, MYH11, NOM1, PRKDC, PSAP, PTDSS1, SERPINC1, SERPINE1, SMPD1, SOD2, UCHL1 | 25 |
| **Familial encephalopathy** | 5.189 | A2M, ALB, ASPM, C3, CDH2, CRYAB, DHCR24, ELOVL1, EPB41L1, FDFT1, FMN2, GALC, GSTM4, H1-4, HIBCH, HMOX1, IRF3, ITIH4, ITM2B, KCNMA1, MECP2, MED17, MMP1, MT-ND5, MT1X, MYH11, NAXE, NDUFB1, NDUFB6, PCLO, PHIP, PSAP, SERPINC1, SERPINE1, SLC16A1, SMPD1, SOD2, TMEM63A, UCHL1, UFM1, UQCR10, XRCC1, YIPF5 | 43 |
| **Progressive encephalopathy** | 5.150 | A2M, ACTC1, ALB, APOC3, C3, CNN1, CRYAB, DHCR24, ELOVL1, F2, FDFT1, FMN2, FUCA1, GALC, HMOX1, HPX, HSPA6, MED17, MMP1, MT-ND5, MYH11, NAXE, NOM1, PIP4P2, PRKDC, PSAP, PTDSS1, SERPINC1, SERPINE1, SMPD1, SOD2, UCHL1 | 32 |
| **Alzheimer disease** | 4.656 | A2M, ALB, APOC3, C3, CNN1, DHCR24, FDFT1, FMN2, HMOX1, HPX, HSPA6, MMP1, MYH11, NOM1, PRKDC, PSAP, PTDSS1, SERPINC1, SERPINE1, SMPD1, SOD2, UCHL1 | 22 |
| **Leukodystrophy** | 4.507 | ALB, GALC, PSAP, SOD2, TMEM63A, UFM1 | 6 |
| **Congenital neurological disorder** | 4.196 | ALB, ASPM, CDH2, DPYSL5, FMN2, FUCA1, GALC, H1-4, HMOX1, MECP2, MED17, MMP1, MT-ND5, MYH11, PCLO, PHIP, PRPS1, PSAP, SERPINC1, SERPINE1, SHOC2, SMPD1, SOD2, YARS2, YIPF5 | 25 |
| **Familial psychiatric disease** | 3.955 | A2M, ALB, ALDH1A3, C3, CRYAB, FDFT1, GALC, HMOX1, ITIH4, ITM2B, KCNMA1, LAMB3, MECP2, MED17, MMP1, MT-ND5, MT1X, NDUFB1, NDUFB6, PHIP, PSAP, SOD2, TMEM63A, UCHL1, UFM1, UQCR10 | 26 |
| **Congenital encephalopathy** | 3.907 | ALB, ASPM, CDH2, DPYSL5, FMN2, GALC, MECP2, MED17, MMP1, MT-ND5, MYH11, PCLO, PHIP, PSAP, SERPINE1, SMPD1, SOD2, YIPF5 | 18 |
| **Leukoencephalopathy** | 3.565 | ALB, GALC, HIBCH, MT-ND5, NAXE, PSAP, SOD2, TMEM63A, UFM1 | 9 |
| **Familial Parkinson disease** | 3.149 | GALC, MT-ND5, PSAP, UCHL1 | 4 |
| **Neuromuscular disease** | 2.971 | A2M, ALB, ATP2C1, BST2, C3, CRYAB, ELOVL1, FDFT1, GALC, HDAC7, HIBCH, HMOX1, ITIH4, KCNMA1, MT-ND5, MT1X, NAXE, NDUFB1, NDUFB6, PIP4P2, PSAP, SGCD, TOP2A, UCHL1, UQCR10 | 25 |
| **Adrenoleukodystrophy** | 2.928 | ALB, SOD2 | 2 |
| **Progressive neuromuscular disease** | 2.807 | ALB, BST2, C3, CRYAB, ELOVL1, FDFT1, GALC, HDAC7, KCNMA1, MT-ND5, PIP4P2, PSAP, TOP2A, UCHL1 | 14 |
| **Progressive motor neuropathy** | 2.337 | ACTC1, ALB, BST2, C3, CRYAB, ELOVL1, FDFT1, FMN2, FUCA1, GALC, HDAC7, KCNMA1, MT-ND5, PIP4P2, PSAP, SOD2, TOP2A, UCHL1 | 18 |
| **Autoimmune neuropathy** | 2.300 | C3, CD59 | 2 |

**Figure 4.**
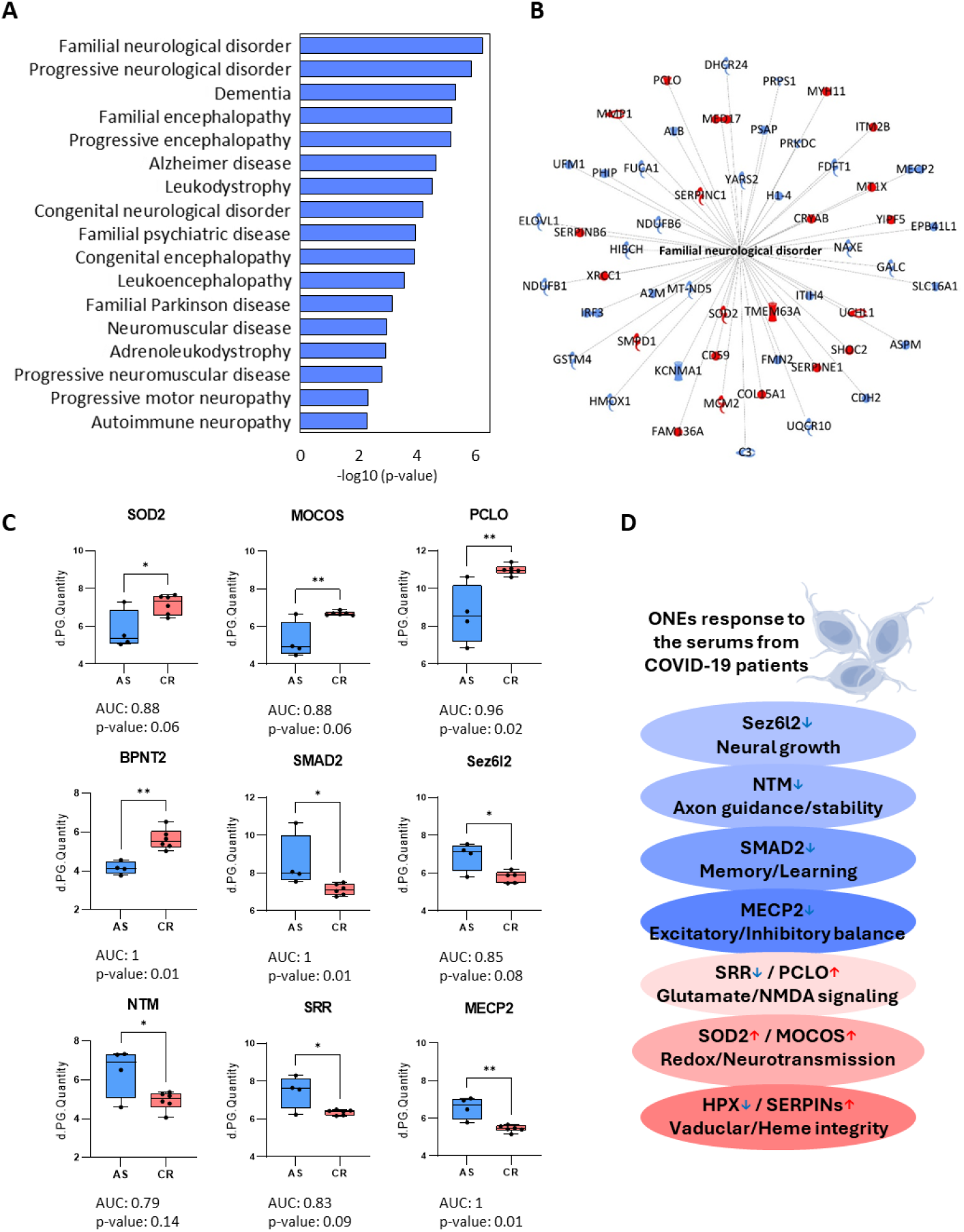
Protein changes associated with neurological and psychiactric disorders. **A)** Functional classification of DEPs associated with neurological disease and psychological disorders. **B)** IPA functional network showing interactions among proteins correlated with familial neurological disorder. **C)** Graphical representation of LFQ intensities for some of the DEPs identified. Data are presented as box-plots, with the median, minimum and maximum values represented, including single data points as well. Differences were considered significant when *p-value < 0.05. **p-value < 0.01. Data from receiver operating characteristic (ROC) analysis with area under curve (AUC) is also indicated. **D**) Summary of proteomic changes in ONEs under COVID-19 stress and their link to long-COVID sequelae.

## 4. DISCUSSION

While SARS-CoV-2 is primarily defined as a respiratory pathogen, a substantial proportion of individuals with Long COVID develop persistent neurological sequelae. These manifestations encompass dysautonomia, particularly orthostatic intolerance, cognitive impairment (“brain fog”), fatigue, headaches, sleep disturbances, and sensory abnormalities such as paresthesia [42]. In addition, psychiatric and neuropsychiatric complications, including anxiety, depression, and delirium, are frequently reported. These symptoms often coexist with sensory deficits, notably ageusia and anosmia, as well as migraines, and in some cases progress to more severe neurological conditions such as encephalopathy or peripheral nervous system disorders [43-45].

Accumulating evidence suggests that the diverse array of neurological conditions associated with long-COVID likely stems from a complex interplay of neuroinflammatory, vascular, metabolic, and neurodegenerative mechanisms [46-49]. However, the precise pathophysiological mechanisms underlying post-COVID neurological dysfunction remain incompletely understood.

In recent years, olfactory neuroepithelium–derived stem cells (ONEs) have been proposed as a translational cellular model for neurological and psychiatric disorders, owing to their unique anatomical and developmental positioning at the interface between the respiratory epithelium and both the peripheral and central nervous systems [50]. However, despite this strategic location, relatively few studies have investigated the specific involvement of ONEs in COVID-19. This is particularly noteworthy given that their anatomical exposure makes them among the first cells to encounter SARS-CoV-2, and that they express key viral entry receptors required for infection. [51, 52]. Briefly, upon entering the nasal cavity, SARS-CoV-2 may interact with olfactory epithelial cells via ACE2 and TMPRSS2 receptors [53], or alternative targets like Neuropilin-1 [54]. The virus may then propagate to the CNS through the olfactory sensory neurons that connect with neurons in the olfactory bulb [17]. Furthermore, evidence from animal models studies suggest that SARS-CoV-2 induces significant damage to the olfactory epithelium, which accounts, among others, for the high prevalence of COVID-19 associated anosmia [55].

To better understand the involvement of ONEs on SARS-CoV-2 entry and the subsequent effects on the nervous system, we employed a LFQ proteomic approach to evaluate the protein changes in ONEs incubated *ex vivo* with the serum from critical (hospitalized) and asymptomatic patients. Our results demonstrate, for the first time to our knowledge, a distinct differential proteomic expression profile in ONEs exposed to the critical patient’s serum (ONEs+CR), compared to those exposed to the asymptomatic-patient serums (ONEs+AS). Furthermore, the changes detected in ONEs+CR cells revealed a dysfunctional pattern associated not only with the viral infective process and inflammation, but also with respiratory and cerebrovascular and nervous system disorders. These findings corroborate the impact that systemic factors from severe COVID-19 infection exert on these cells.

Among the proteins directly correlated with COVID-19 infection, matrix metalloproteinase-1 (MMP1), a collagenase that acts as an agonist for protease activated receptor 1 (PAR1) across various cell types, was found up-regulated in ONEs+CR cells. In the context of the nervous system, the MMP1-PAR1 axis can significantly affect proliferation and differentiation of neural progenitor cells [56, 57]. Furthermore, increased MMP1 might contribute to neuronal damage associated with degenerative and inflammatory conditions [58]. Thus, up-regulation of MMP1 in ONEs+CR cells suggests that the serum of critical patients may induce a pro-inflammatory signaling cascade that compromises neural integrity at the olfactory interface. Notably, elevated serum levels of MMP1 have been observed in COVID-19 patients in correlation with disease severity [59], and several MMP1 inhibitors are currently being investigated as potential therapeutics to mitigate the inflammatory response to SARS-CoV-2 [60].

The lysosome-associated membrane glycoprotein 2 (LAMP2) was also significantly up-regulated in ONEs+CR cells. This protein has been previously linked to chaperone-mediated autophagy [61], a process that appears dysregulated during COVID-19 infection [62]. Of interest, LAMP2 upregulation was also observed in a similar study, exposing endothelial colony forming cells (ECFCs) to the serum factors of critical COVID19 patients [26].

Other up-regulated proteins in ONEs+CR cells associated to COVID19 infection and the risk of thrombosis and ischemia include two members of the serpin family: SERPINC1 and SERPINE1. These proteins might have an important role in SARS-CoV-2 infection, given the role of SERPINC1 as an anticoagulant factor, or the inhibitory effect of SERPINE1 over plasminogen activator. Interestingly, both SERPINC1 and SERPINE1 have been shown to prevent infection by TMPRSS2-mediated S-protein cleavage, thereby blocking viral entry [63]. Conversely, other proteins related to vascular protection and heme homeostasis, such as Heme oxygenase 1 (HMOX1) and Hemopexin (HPX), were downregulated in ONEs+CR cells. These proteins are involved in heme degradation, which is also associated with inflammation and coagulation, and therefore with thrombosis [64]. HPX binds free heme and maintains red blood cells (RBCs) integrity [64, 65]; notably, RBCs deformability has been linked to inflammatory conditions and hypoxia in COVID-19 patients, contributing to cerebrovascular complications [65]. Both HMOX1 and HPX have been described as markers of COVID-19 severity [64, 66, 67]. Similarly, inter α-trypsin inhibitor heavy chain 4 (ITIH4) was also down-regulated in ONEs pre-stimulated with critical-patients serum. ITIH4 is a protease inhibitor previously described by proteomics as being altered in the serum of COVID-19 patients, alongside other members of the inter-α-trypsin inhibitors family [68, 69], such as the ITIH3 protein, which was also down-regulated in our ONEs+CR cells.

In terms of neurological and psychiatric disorders, previous studies have reported that the most common sequelae following SARS-CoV2 infection are anxiety, depression, post-traumatic symptoms, cognitive impairment, and sleep disturbance [70, 71]. Remarkably, many of the DEPs identified in ONEs+CR compared to ONEs+AS, correlated with dementia, encephalopathy, leukodystrophy, neuromuscular diseases or familiar psychiatric disease, among others. Among the proteins up-regulated in ONEs+CR we identified the proteins like Superoxide Dismutase 2 (SOD2) and molybdenum cofactor sulfurase (MOCOS), both associated with redox imbalance and neurodegenerative diseases. Indeed, SOD2 has been previously described as altered in COVID-19 patients in association with increased oxidative stress [72], and it seems implicated in the progression of neurodegenerative diseases such as Alzheimer and Parkinson, or cerebrovascular stroke [33]. Furthermore, MOCOS has been identified as altered in nasal olfactory stem cells of adults with autism spectrum disorders [34, 35], and it has been postulated to play a critical role in redox homeostasis and neurotransmission [34, 73]. To our knowledge, this is the first report linking altered MOCOS expression to COVID-19 pathology. Likewise, the protein bisphosphate nucleotidase 2 (BPNT2) was also up-regulated in ONEs incubated with the serum from critical patients. BPNT2 is a known target inhibited by lithium, one of the most widely used treatment for mental illnesses such as bipolar disorder and depression. Furthermore, lithium interferes in immune and inflammatory responses, and aids in the normalization of cytokine levels [38, 39]. Consequently, several studies investigating lithium as a COVID-19 therapeutic have shown that it reduces progression to critical states in COVID-19 patients and decreases long-term adverse outcomes [39].

Conversely, among the proteins down-regulated in ONEs+CR, we identified several proteins associated with neurogenesis and neuronal function: seizure-related 6 homolog like 2 (Sez6l2), Neurotrimin (NMT), SMAD2 and methyl CpG binding protein 2 (MECP2). Sez6l2 is highly expressed in the hippocampus and cerebellar cortex, where it contributes to neuronal growth. Indeed, Sez6l2 knockdown increased neurite outgrowth and modulated neuronal differentiation [41, 74]. Moreover, Sez6l2 dysregulation has been linked to autism spectrum disorder and other psychiatric disorders, including depression, schizophrenia, and bipolar disorder [74, 75]. Similarly, the transcription factor SMAD2 is also involved in neurodevelopment, influencing cell fate determination and the formation of dendrites, axons, and synapses [40]. Given that SMAD2 has recently been shown to affect hippocampal-related tasks such as long-term learning and working memory [40], its reduction following severe COVID-19 may contribute to the memory deficits and brain fog reported by Long-COVID patients. Furthermore, NTM, a neural cell adhesion molecule, promotes axonal fasciculation, axonal guidance and synaptic stabilization [76]. Its loss in the ONE+CR, suggests a disruption in the structural integrity of the neural circuit, potentially contributing to the prolonged sensory and cognitive disturbances observed in the chronic phase of the disease. Finally, MECP2 is involved in several stages of neurodevelopment, from prenatal neurogenesis to postnatal synaptic connections and function. Loss of function or mutations leads to major neurodevelopmental disorders such as Rett syndrome, characterized by delayed cognitive and motor development and impaired social interaction [77]. In addition, MECP2 deficiency may disrupt the balance between excitatory and inhibitory neuronal circuits, particularly when MECP2 loss occurs in inhibitory neurons [77]. Therefore, MECP2 downregulation in ONE+CR might be associated with cognitive difficulties often associated with Long-COVID sequelae. Overall, the simultaneous down-regulation of these proteins suggest that the systemic environment of COVID-19 critical patients, represented in their serum factors, may negatively affect neural plasticity, providing a compelling molecular framework for the persistent cognitive and sensory deficits characteristics of Long-COVID. Further studies are required to confirm our findings.

Although psychotic symptoms are not among the most commonly reported psychiatric manifestations of long COVID-19 [78], growing evidence suggests that SARS-CoV-2 infection may act as a biological stressor capable of increasing vulnerability to psychosis, including schizophrenia-spectrum disorders [79]. In line with this possibility, our findings show that the expression of serine racemase (SRR), a gene implicated in schizophrenia risk, was significantly reduced in ONEs following incubation with CR serum. SRR catalyzes the conversion of L-serine to D-serine, an essential co-agonist of glutamatergic NMDA receptors, whose hypofunction has been strongly associated with the pathophysiology of schizophrenia [36]. Indeed, reduced brain levels of SRR have been reported in patients with schizophrenia [80], and SRR knockout mice exhibited features associated with this mental disorder, including cognitive impairment, increased lateral ventricle size, cortical atrophy, reduced dendritic length and density, and downregulation of the cortical GABAergic interneurons [80]. Additionally, the piccolo presynaptic cytomatrix protein (PCLO) was up-regulated in ONEs+CR. Interestingly, PCLO is also associated with the NMDA receptor co-agonist, D-serine. It has been described that increased PCLO levels may serve as a homeostatic response of dopaminergic system to attenuate excessive dopaminergic synaptic plasticity [81]. Furthermore, dysregulation of this protein may be involved in the presynaptic matrix pathogenesis of affective disorders and schizophrenia [36]. Indeed, PCLO polymorphisms have been described in major depression and bipolar disorder, with specific mutations also documented in a case of schizophrenia [36, 37]. Taken together, the proteomic alterations observed in ONEs likely have an impact on neural function, although further studies are required to link these proteins to neurological or psychiatric symptoms observed in COVID-19 patients.

## Conclusion

In summary, this study provides the first proteomic characterization of ONEs exposed to the systemic environment of critical COVID-19 patients. Our findings demonstrate that serum factors from severe cases trigger a profound proteomic shift in neural progenitors, characterized by the up-regulation of oxidative stress markers, such as SOD2 or MOCOS, and the down-regulation of essential neuroplasticity drivers. Moreover, the simultaneous disruption of NMDA receptor modulation, via SRR and PCLO, and the loss of excitatory-inhibitory balance, via MECP2, may be representative of the molecular underpinnings of the neurological disturbances and psychiatric sequelae that define Long-COVID. While further research with a higher sample size is necessary to validate our findings, and determine their relevance across diverse biological backgrounds, the *ex-vivo* approach described here constitutes an optimal model for studying the response of ONEs to SARS-CoV-2 infection, and a promising platform for investigating PASC. Ultimately, this study reveals new proteomic targets that may provide critical insights into the neurobiological complications and long-term adverse outcomes in post-COVID-19 patients.

