## Supplementary Table 1 for "The serum from critical COVID-19 patients induces proteomic changes in olfactory neuroepithelial cells that resemble post-covid neurological complications"

### Supplementary material

**Supplementary table S1. Quantitative analysis of proteins differentially expressed in ONEs after COVID-19 patient's serum incubation (ONEs CR*vs*AS).** The table includes (from left to right): Protein ID (Uniprot accession number), gen name, CR*vs*AS difference and significance (-$\log_{10}$ p-value). The table shows the significant values (p-value < 0.05) over-expressed (difference > 1; red) or under-expressed (difference < 1; green).

| Protein ID | Gen name | CR*vs*AS difference | -$\log_{10}$ (p-value) |
|---|---|---|---|
| **P31513** | FMO3 | 2.7484 | 2.4710 |
| **P52815** | MRPL12 | 2.4763 | 2.5702 |
| **Q9ULV0** | MYO5B | 2.3636 | 2.2525 |
| **Q9Y6V0** | PCLO | 2.3484 | 2.2306 |
| **P63267** | ACTG2 | 2.3355 | 2.2291 |
| **P17405** | SMPD1 | 2.2816 | 2.4899 |
| **Q6AWC2** | WWC2 | 2.2526 | 2.1037 |
| **P35237** | SERPINB6 | 2.1756 | 2.0924 |
| **P48651** | PTDSS1 | 2.1593 | 1.7860 |
| **P51397** | DAP | 2.1254 | 2.1054 |
| **P62861** | FAU | 2.0487 | 1.6432 |
| **Q9UQ13** | SHOC2 | 2.0429 | 2.0465 |
| **Q8NBT0** | POC1A | 2.0191 | 2.5895 |
| **P42126** | ECI1 | 2.0175 | 2.0818 |
| **P03956** | MMP1 | 1.9055 | 1.9013 |
| **Q10589** | BST2 | 1.8568 | 2.3059 |
| **Q969M3** | YIPF5 | 1.8476 | 2.3743 |
| **P28300** | LOX | 1.7788 | 2.4472 |
| **P02511** | CRYAB | 1.7728 | 2.3201 |
| **Q9BPZ7** | MAPKAP1 | 1.7298 | 2.3548 |
| **P30711** | GSTT1 | 1.7284 | 2.0030 |
| **Q8IVF2** | AHNAK2 | 1.7086 | 1.9283 |
| **Q8NFU3** | TSTD1 | 1.6951 | 1.3569 |
| **Q9UBB5** | MBD2 | 1.6728 | 2.1371 |
| **Q9NVC6** | MED17 | 1.5875 | 1.6774 |
| **Q03395** | ROM1 | 1.5654 | 2.1654 |
| **P98179** | RBM3 | 1.5586 | 2.7733 |
| **Q9H0J9** | PARP12 | 1.5289 | 1.8612 |
| **Q96C01** | FAM136A | 1.4948 | 1.6150 |
| **P62312** | LSM6 | 1.4872 | 2.9072 |
| **Q9NX62** | BPNT2 | 1.4843 | 2.9459 |
| **Q96EN8** | MOCOS | 1.4566 | 2.2441 |
| **Q96ET8; Q9NYZ1** | TVP23C; TVP23B | 1.4397 | 2.0325 |
| **Q8WXC6** | COPS9 | 1.4134 | 2.3789 |
| **P18887** | XRCC1 | 1.4025 | 2.3814 |
| **P04179** | SOD2 | 1.4007 | 1.6742 |
| **Q9BTZ2** | DHRS4 | 1.3692 | 2.0151 |
| **P68032** | ACTC1 | 1.3635 | 2.2028 |
| **Q8WUI4** | HDAC7 | 1.3387 | 1.4536 |
| **O76011** | KRT34 | 1.3191 | 5.8349 |

| | | | |
|---|---|---|---|
| **P50238** | CRIP1 | 1.3079 | 2.0594 |
| **Q9NPJ3** | ACOT13 | 1.2953 | 1.8340 |
| **O14558** | HSPB6 | 1.2835 | 2.1397 |
| **P25208** | NFYB | 1.2651 | 2.2286 |
| **P51911** | CNN1 | 1.2452 | 2.3669 |
| **Q99541** | PLIN2 | 1.2359 | 1.4426 |
| **P56556** | NDUFA6 | 1.2188 | 1.8576 |
| **P39059** | COL15A1 | 1.2178 | 2.6037 |
| **P13489** | RNH1 | 1.2093 | 2.3342 |
| **P13473** | LAMP2 | 1.2082 | 2.5870 |
| **Q6NUQ1** | RINT1 | 1.2081 | 3.1423 |
| **Q96NT0** | CCDC115 | 1.1963 | 1.3056 |
| **Q9UNN8** | PROCR | 1.1944 | 1.6978 |
| **P0CW19; P0CW20** | LIMS3; LIMS4 | 1.1922 | 1.6097 |
| **Q9NRX2** | MRPL17 | 1.1890 | 1.6131 |
| **P14406** | COX7A2 | 1.1883 | 2.6423 |
| **P54819** | AK2 | 1.1859 | 2.9476 |
| **P48509** | CD151 | 1.1745 | 1.4476 |
| **Q9UIC8** | LCMT1 | 1.1726 | 2.0735 |
| **Q96C19** | EFHD2 | 1.1607 | 2.6046 |
| **P50453** | SERPINB9 | 1.1449 | 2.1907 |
| **Q15121** | PEA15 | 1.1394 | 2.6628 |
| **Q92629** | SGCD | 1.1089 | 2.6909 |
| **Q4L180** | FILIP1L | 1.1056 | 2.0392 |
| **O94886** | TMEM63A | 1.1026 | 1.6824 |
| **Q96RQ3** | MCCC1 | 1.0968 | 1.8310 |
| **Q04828** | AKR1C1 | 1.0923 | 2.6229 |
| **P80297** | MT1X | 1.0872 | 1.7954 |
| **P47712** | PLA2G4A | 1.0689 | 1.9300 |
| **P05121** | SERPINE1 | 1.0684 | 1.4903 |
| **P62308** | SNRPG | 1.0602 | 1.5503 |
| **P01008** | SERPINC1 | 1.0561 | 3.4828 |
| **P35749** | MYH11 | 1.0556 | 1.5198 |
| **P47895** | ALDH1A3 | 1.0534 | 2.2628 |
| **O95456** | PSMG1 | 1.0495 | 1.8010 |
| **P07951** | TPM2 | 1.0469 | 2.5340 |
| **P49736** | MCM2 | 1.0401 | 1.8118 |
| **P13987** | CD59 | 1.0389 | 3.3496 |
| **P09234** | SNRPC | 1.0339 | 1.9860 |
| **P09936** | UCHL1 | 1.0315 | 2.7041 |
| **P98194** | ATP2C1 | 1.0308 | 3.7813 |
| **Q8ND56** | LSM14A | 1.0265 | 2.5018 |
| **P17661** | DES | 1.0254 | 1.3034 |
| **Q9Y287** | ITM2B | 1.0157 | 1.4757 |
| **P62995** | TRA2B | 1.0038 | 2.8052 |
| **O00425** | IGF2BP3 | 1.0013 | 2.1212 |
| **Q9UBU6** | FAM8A1 | -1.0025 | 1.9511 |
| **P43235** | CTSK | -1.0069 | 2.2156 |

| | | | |
|---|---|---|---|
| **Q9Y6U3** | SCIN | -1.0096 | 1.8057 |
| **P61960** | UFM1 | -1.0101 | 2.6298 |
| **Q9NP58** | ABCB6 | -1.0141 | 1.7647 |
| **O60565** | GREM1 | -1.0186 | 2.7487 |
| **Q8N4L2** | PIP4P2 | -1.0224 | 1.9754 |
| **P04066** | FUCA1 | -1.0247 | 1.9221 |
| **Q9BXP2** | SLC12A9 | -1.0266 | 1.4606 |
| **Q5C9Z4** | NOM1 | -1.0379 | 2.1175 |
| **P17066** | HSPA6 | -1.0404 | 1.4916 |
| **Q8WWQ0** | PHIP | -1.0426 | 1.7153 |
| **P53985** | SLC16A1 | -1.0443 | 2.6907 |
| **Q9NP90** | RAB9B | -1.0460 | 2.1997 |
| **A6NIH7** | UNC119B | -1.0480 | 1.8879 |
| **O14763** | TNFRSF10B | -1.0509 | 2.5685 |
| **Q15124** | PGM5 | -1.0549 | 2.2456 |
| **Q6NVY1** | HIBCH | -1.0552 | 2.6121 |
| **O14531** | DPYSL4 | -1.0568 | 1.5983 |
| **P11388** | TOP2A | -1.0616 | 1.6299 |
| **Q8NCW5** | NAXE | -1.0647 | 1.6451 |
| **P0CG30** | GSTT2B | -1.0713 | 2.3769 |
| **Q9BSA9** | TMEM175 | -1.0725 | 1.7502 |
| **P04004** | VTN | -1.0785 | 1.4484 |
| **P56134** | ATP5MF | -1.0799 | 2.0260 |
| **Q9GZT4** | SRR | -1.0853 | 1.8239 |
| **P60059** | SEC61G | -1.0907 | 1.8677 |
| **Q9BRK3** | MXRA8 | -1.0924 | 1.7118 |
| **Q9BW60** | ELOVL1 | -1.0932 | 3.2697 |
| **A5PLL7** | PEDS1 | -1.0948 | 2.1586 |
| **Q9UDW1** | UQCR10 | -1.0950 | 1.7403 |
| **Q9NZ56** | FMN2 | -1.0997 | 1.4512 |
| **Q9UKW4** | VAV3 | -1.1003 | 2.3431 |
| **Q6UXD5** | SEZ6L2 | -1.1134 | 1.7128 |
| **P54803** | GALC | -1.1135 | 2.5083 |
| **P51608** | MECP2 | -1.1159 | 2.6995 |
| **Q8TDB4** | MGARP | -1.1284 | 1.8802 |
| **P02768** | ALB | -1.1318 | 1.8881 |
| **Q8NHQ8** | RASSF8 | -1.1365 | 1.4157 |
| **Q4KMZ1** | IQCC | -1.1428 | 1.9279 |
| **O95139** | NDUFB6 | -1.1468 | 1.9681 |
| **Q8N5W9** | RFLNB | -1.1535 | 2.7120 |
| **P02790** | HPX | -1.1554 | 4.0125 |
| **Q01831** | XPC | -1.1561 | 1.8224 |
| **Q9C075** | KRT23 | -1.1578 | 2.5193 |
| **P37268** | FDFT1 | -1.1590 | 2.0578 |
| **Q6NUK4** | REEP3 | -1.1607 | 2.9079 |
| **O75438** | NDUFB1 | -1.1609 | 1.5284 |
| **Q53LP3** | SOWAHC | -1.1683 | 2.6847 |
| **P78527** | PRKDC | -1.1718 | 2.3027 |

| | | | |
|---|---|---|---|
| **Q96B49** | TOMM6 | -1.1774 | 1.8612 |
| **Q9NZQ3** | NCKIPSD | -1.1811 | 3.2848 |
| **Q9Y679** | AUP1 | -1.1891 | 2.5919 |
| **Q96HD1** | CRELD1 | -1.1914 | 1.8821 |
| **Q8IVF7** | FMNL3 | -1.1934 | 1.4134 |
| **P07602** | PSAP | -1.2046 | 2.0314 |
| **Q13308** | PTK7 | -1.2077 | 2.2369 |
| **P09601** | HMOX1 | -1.2110 | 2.0860 |
| **Q5T9A4** | ATAD3B | -1.2129 | 2.1512 |
| **Q9Y2Z4** | YARS2 | -1.2168 | 2.0742 |
| **Q8N3E9** | PLCD3 | -1.2237 | 1.9540 |
| **P10412** | H1-4 | -1.2239 | 1.3726 |
| **P83731** | RPL24 | -1.2347 | 2.4091 |
| **Q9NP92** | MRPS30 | -1.2365 | 1.4516 |
| **Q93034** | CUL5 | -1.2584 | 2.7474 |
| **P16403** | H1-2 | -1.2815 | 2.1791 |
| **Q12791** | KCNMA1 | -1.2931 | 2.2745 |
| **P62805** | H4C1 | -1.3005 | 1.7415 |
| **Q9Y3C0** | WASHC3 | -1.3164 | 1.3630 |
| **Q8TAD7** | OCC1 | -1.3203 | 2.0993 |
| **Q9NVM1** | EVA1B | -1.3430 | 2.5169 |
| **Q5SNT2** | TMEM201 | -1.3609 | 1.3853 |
| **P16234** | PDGFRA | -1.3743 | 2.1304 |
| **Q9H2H8** | PPIL3 | -1.3832 | 2.1480 |
| **Q6T4R5** | NHS | -1.4243 | 1.3878 |
| **Q15796** | SMAD2 | -1.4271 | 1.4055 |
| **P03915** | MT-ND5 | -1.4544 | 1.5953 |
| **Q658P3** | STEAP3 | -1.4999 | 2.3288 |
| **Q9P121** | NTM | -1.5108 | 1.5553 |
| **Q9BPU6** | DPYSL5 | -1.5140 | 2.6486 |
| **P60891** | PRPS1 | -1.5201 | 2.1418 |
| **Q8TCZ2** | CD99L2 | -1.5226 | 2.0405 |
| **Q99424** | ACOX2 | -1.5314 | 2.3225 |
| **Q9Y547** | HSPB11 | -1.5571 | 1.8221 |
| **P00734** | F2 | -1.6071 | 2.5482 |
| **Q9Y2J8** | PADI2 | -1.6373 | 2.1472 |
| **P69905** | HBA1 | -1.6393 | 2.0999 |
| **Q8ND90** | PNMA1 | -1.6578 | 2.0951 |
| **Q4V9L6** | TMEM119 | -1.6630 | 1.6957 |
| **Q9Y6X4** | FAM169A | -1.6966 | 2.2596 |
| **O15083** | ERC2 | -1.7036 | 2.1259 |
| **P01024** | C3 | -1.7140 | 2.4186 |
| **Q06033** | ITIH3 | -1.7150 | 2.4750 |
| **Q03013** | GSTM4 | -1.7556 | 2.0441 |
| **P01023** | A2M | -1.7899 | 4.8596 |
| **Q14653** | IRF3 | -1.8417 | 1.6222 |
| **P16401** | H1-5 | -1.8740 | 1.8569 |
| **Q15392** | DHCR24 | -1.8920 | 2.7228 |

| | | | |
|---|---|---|---|
| **Q9BRX8** | PRXL2A | -1.9222 | 1.7011 |
| **O14495** | PLPP3 | -1.9588 | 1.9915 |
| **P22676** | CALB2 | -1.9668 | 2.8177 |
| **P19022** | CDH2 | -1.9885 | 1.3498 |
| **Q9UKG9** | CROT | -2.0161 | 2.3016 |
| **P02656** | APOC3 | -2.0271 | 2.4177 |
| **P84243** | H3-3A | -2.0433 | 2.5004 |
| **Q8WUN7; Q9HAC8** | UBTD2; UBTD1 | -2.0734 | 1.9594 |
| **Q14624** | ITIH4 | -2.1376 | 3.3102 |
| **Q9H4G0** | EPB41L1 | -2.2397 | 2.1418 |
| **Q14534** | SQLE | -2.3955 | 1.5518 |
| **Q9BS34** | ZNF670 | -2.4533 | 4.1337 |
| **Q8IZT6** | ASPM | -2.8330 | 2.8530 |
| **Q9BW72** | HIGD2A | -2.8405 | 1.7519 |
| **Q13751** | LAMB3 | -2.8963 | 1.6222 |
